# Loss of MLKL Decreases Necrotic Core but Increases Macrophage Lipid Accumulation In Atherosclerosis

**DOI:** 10.1101/746644

**Authors:** Adil Rasheed, Sabrina Robichaud, My-Anh Nguyen, Michele Geoffrion, Mary Lynn Cottee, Taylor Dennison, Antonietta Pietrangelo, Richard Lee, Mireille Ouimet, Katey J Rayner

**Author notes:** Please address correspondence to: Katey J Rayner, University of Ottawa Heart Institute, 40 Ruskin Street, Room H4211A, Ottawa, Ontario, Canada, K1Y 4W7.

## Abstract

**Objectives:** During the advancement of atherosclerosis, the cellularity of the plaque is governed by the influx of monocyte-derived macrophages and their turnover via apoptotic and non-apoptotic forms of cell death. Previous reports have demonstrated that programmed necrosis, or necroptosis, of macrophages within the plaque contribute to necrotic core formation. Knockdown or inhibition of the necrosome components RIPK1 and RIPK3 slow the progression of atherosclerosis, and activation of the terminal step of necroptosis, MLKL, has been demonstrated in advanced human atherosclerotic plaques. However, whether MLKL directly contributes to lesion development and necrotic core formation has not been investigated.

**Approaches and Results:** MLKL expression was knocked down in atherogenic *Apoe*- knockout mice via subcutaneous administration of antisense oligonucleotides (ASO). During advanced atherogenesis, *Mlkl* knockdown potently reduced cell death in the plaque, with a significant reduction in the necrotic core. However, total lesion area in the aortic sinus remained unchanged. Furthermore, treatment with the MLKL ASO unexpectedly reduced circulating cholesterol levels compared to control ASO, while staining for lipids within the plaque was significantly increased. Peritoneal macrophages transfected with the MLKL ASO showed increased lipid loading upon incubation with modified cholesterol-rich lipoproteins. In lipid-loaded macrophages, MLKL co-localized with Rab7, a marker of the late endosome.

**Conclusions:** These studies confirm the requirement for MLKL as the executioner of necroptosis, and as such a significant contributor to the necrotic core during atherogenesis. We also identified a previously unknown role for MLKL in interacting with endosomal trafficking components to regulate lipid uptake in macrophages during atherogenesis.

## INTRODUCTION

The development of atherosclerosis occurs throughout life, but is only diagnosed after the growing atheroma impedes arterial blood flow^1^. Initially, plaques are lipid-rich and relatively benign, but during the advanced stages, the composition of the plaque changes and pro-inflammatory processes increase lesion vulnerability. A vulnerable plaque is more prone to rupture, resulting in myocardial infarction or stroke, which account for approximately 75% of cardiovascular disease deaths^1, 2^. During these later stages of atherosclerosis, cellular turnover in the plaque becomes dysregulated as macrophage foam cells undergo both apoptotic and non-apoptotic modes of cell death but are inefficiently cleared, all of which contribute to the acellular necrotic core^3^. When dead cell membranes rupture, there is a release of the intracellular contents that serve as damage-associated molecular patterns (DAMPs)^4^. Therefore, these dying macrophages contribute to late stage pathogenesis by i) the release of intracellular pro-inflammatory mediators and ii) the production of the necrotic core that promotes plaque vulnerability and rupture.

Traditionally, the necrotic debris found in lesions was believed to be caused by passive lysis of remnant apoptotic cells, but a recently described mode of cell death, known as necroptosis has been shown to contribute to this so-called ‘cellular explosion’^5^. Necrotic and necroptotic cells share similar morphology, including cellular blebbing and membrane rupture, however unlike necrosis, necroptosis is programmed and dictated by a signaling pathway^5^. Engagement of death receptors, such as TNF and Toll-like receptors at the cell surface promotes the sequential phosphorylation of the receptor interacting protein kinases 1 and 3 (RIPK1 and RIPK3)^5^. This necrosome complex then recruits and phosphorylates the mixed lineage kinase domain-like protein (MLKL). Activated MLKL homo-oligomerizes and localizes to the membrane to facilitate cell lysis^6, 7^. MLKL was originally discovered as the executioner of necroptosis via siRNA and small molecule screens of necrotic cell death inhibitors^8, 9^. This was later confirmed in *Mlkl*-knockout mice which were resistant to necroptotic death induced by LPS or TNFα and the pan-caspase inhibitor zVAD.fmk *in vitro* or experimental pancreatitis *in vivo*^10^. MLKL at its discovery was characterized as a pseudokinase, having no intrinsic kinase capability but must itself be phosphorylated to execute necroptosis^11^. More recently, additional non-necroptotic roles of MLKL have been uncovered, including activation of the inflammasome^12, 13^ and binding to ESCRT proteins during extracellular vesicle secretion^14^. Thus, although MLKL is unequivocally involved in necroptotic cell death, its other roles in the cell have not yet been fully explored.

We and others have shown that inhibition of necroptosis via blocking RIPK1 kinase activity or genetic deletion of RIPK3 in atherosclerotic mouse models potently reduce both necrotic core and total plaque area in the aortic sinus^15, 16^. Our group has also identified necroptosis as an active process in human atherosclerosis, where phosphorylated MLKL was detected in late stage coronary atherosclerotic plaques but not early atheromas^15^. However, whether MLKL is a direct contributor to atherosclerosis has not been previously explored. Here we set out to investigate whether inhibition of MLKL decreases the advancement of the atherosclerotic plaque. *Mlkl* expression was systemically knocked down by administration of antisense oligonucleotides (ASOs) in *Apoe*-knockout mice. Consistent with its known function in necroptosis, inhibition of *Mlkl* reduced necrotic core size and decreased cell death in late-stage plaques and in cultured macrophages. Unexpectedly, this did not lead to a change in total plaque area, despite reductions in circulating cholesterol levels. Furthermore, *Mlkl* knockdown significantly increased the lipid content of the atherosclerotic plaque. Macrophages receiving the MLKL ASO showed a significant increase in lipid droplets when incubated with atherogenic lipoproteins, and in macrophages loaded with oxidized or aggregated LDL, we found colocalization of MLKL within the late endosome. These data therefore indicate that beyond promoting macrophage necroptosis, MLKL also plays an anti-atherogenic role in repressing lipid accumulation within plaque macrophages.

## MATERIALS AND METHODS

Material and Methods are available in the online-only Data Supplement.

## RESULTS

### Mlkl knockdown decreases necrotic core in the atherosclerotic plaque

Previous reports have shown that inhibition of necroptotic pathway components RIPK1 and RIPK3 reduce the necrotic core area of the atherosclerotic plaques^15, 16^. We thus reasoned that inhibition of MLKL, which is downstream of RIPK1 and RIPK3 and is the committed step within the necroptosis pathway, would similarly reduce the necrotic core of the plaque. To this end, we administered two independent sequences of ASOs targeting MLKL to *Apoe*-knockout (*Apoe*-/-) mice. Knockdown of MLKL was confirmed at both the gene and protein expression levels in the livers of *Apoe*-/- mice after 8 weeks of ASO treatment (**Supplemental Figure 1**). While necrotic core was not altered during early atherogenesis (**Figure 1A-B**), *Mlkl* knockdown reduced the necrotic core area by >50% during late stage atherosclerosis (**Figure 1C**). Despite this reduction in necrotic core, and unlike inhibition of RIPK1 & 3, silencing MLKL did not change total lesion area (**Figure 1D**). Surprisingly, we observed a reduction in circulating cholesterol levels at both 8 and 16 weeks of Western diet feeding in MLKL ASO treated mice (21-29%; *p*<0.0001, *p*<0.05; **Figure 1E**). These reductions in circulating cholesterol were present in all lipoprotein fractions (**Supplemental Figure 2**). To confirm that these observations were not specific for this particular MLKL targeting sequence, we also examined the development of atherosclerosis with a distinct MLKL-targeted ASO and observed the same trends in plaque area and circulating cholesterol (**Supplemental Figure 2A-D**). Apart from its role as the executioner of necroptosis, MLKL has been demonstrated to activate the inflammasome in bone marrow derived macrophages and within the intestinal epithelium^12, 13^. To determine whether *Mlkl* knockdown altered the inflammatory status of the recipient *Apoe*-/- mice, we assessed surrogate markers of systemic inflammation. There were no changes in the complete blood counts of circulating inflammatory cells known to contribute to atherosclerosis at either 8 weeks or 16 weeks of treatment (**Supplemental Figure 2E-F**). Furthermore, ELISA analysis of the serum at 16 weeks did not reveal changes in circulating pro-inflammatory cytokines IL-1β, IL-6, MCP-1, or CCL-5 (**Supplemental Figure 2G**), together suggesting that treatment with the MLKL ASO did not significantly change systemic inflammation to drive atherogenesis. Therefore, although MLKL knockdown reduced necrotic cell death in the plaque, the discrepancy between necrotic core and total plaque area suggested that systemic knockdown of *Mlkl* may also elicit non-necroptotic functions that drive atherogenesis.

**Figure 1.**
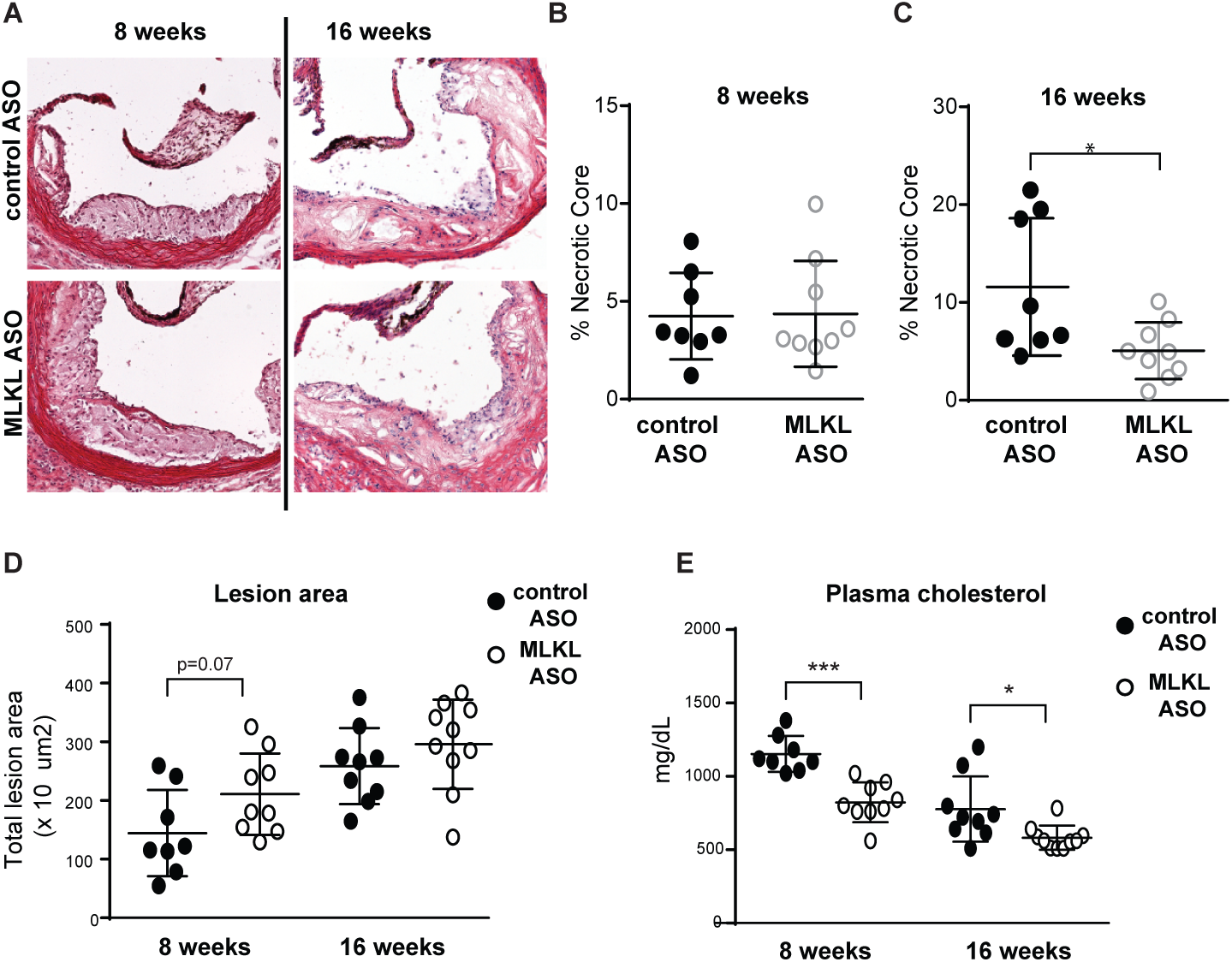
*Mlkl* knockdown by administration of antisense oligonucleotides to *Apoe*-knockout decreases necrotic core but not total plaque area. *Apoe*-knockout (*Apoe*-/-) mice were fed a Western diet for 8 or 16 weeks while receiving control or MLKL ASO. **A**, Aortic sinus sections were stained with hematoxylin and eosin. Scale bar=200μm. The necrotic core of aortic sinus plaques were quantified at **B**, 8 and **C**, 16 weeks of treatment. **D**, Total plaque area in the aortic sinus and **E**, cholesterol levels in the serum were evaluated at both the 8 and 16 week timepoints. n=8-10 mice per group. Data for MLKL ASO Sequence 2 can be found in Supplemental Figures 1-2. Complete blood counts were performed in whole blood after **F**, 8 and **G**, 16 weeks of treatment. **H**, Circulating pro-inflammatory cytokines were quantified by multiplex ELISA in the serum of *Apoe*-/- mice treated for 16 weeks (select factors shown). n=6-11 mice per group. Data presented as mean±SD. \**P*<0.05, \*\*\*\**P*<0.0001.

**Figure 2.**
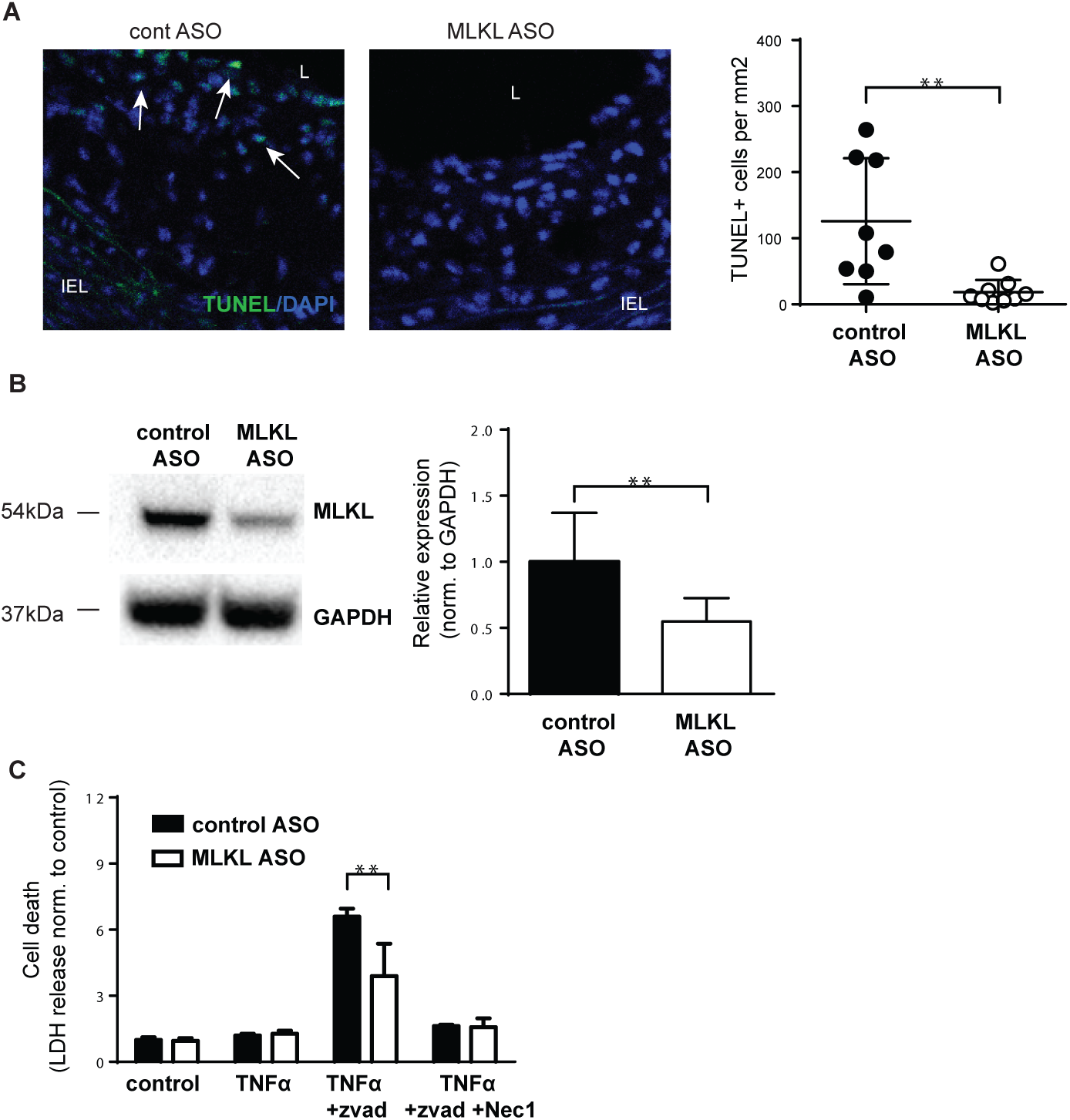
MLKL ASO decreases cell death in the plaque and in cultured macrophages. **A**, TUNEL staining was performed in aortic sinus plaques from *Apoe*-/- mice receiving ASOs for 8 weeks. Scale bar=100μm. Data presented as mean±SD. n=8-9 mice per group. **B**, *Mlkl* knockdown after ASO treatment transfected peritoneal macrophages was quantified by Western blotting. **C**, Macrophage cell death was assessed in response to TNFα (50ng/ml), zVAD.fmk (zVAD, 50μM), and Necrostatin-1 (Nec-1, 50μM). Data presented as mean±SEM. Representative experiment using n=6- 11 mice per group. \*\**P*<0.01.

### MLKL drives necroptotic cell death in the atherosclerotic plaque

To confirm that the reduction in necrotic core formation in the atherosclerotic plaque was due to a reduction in cell death in the absence of MLKL, we performed TUNEL staining to positively identify breaks in the DNA typically found in the late phases of cell death. We observed a striking reduction in TUNEL staining in the plaques of MLKL ASO treated mice, prior to the development of the necrotic core (**Figure 2A**). To confirm that *Mlkl* knockdown by ASO does indeed decrease necroptotic cell death, peritoneal macrophages from WT mice were transfected with control or MLKL ASO and cell death was analyzed in response to prototypical necroptotic ligands. We first confirmed Mlkl was reduced in MLKL ASO treated macrophages by immunoblotting (**Figure 2B**). Cell death of the macrophages was then assessed in response to co-treatment with TNFα, to engage the death receptors, and zVAD.fmk, a pharmacologic pan-caspase inhibitor to push cell death towards necroptosis. Macrophages transfected with MLKL ASO showed significantly decreased TNFα+zVAD.fmk induced cell death as compared to control ASO (**Figure 2C**). This cell death was necroptotic specific as cell death was inhibited by treatment with Necrostatin-1, a small molecule inhibitor of the RIPK1 & 3 signaling^17^. These data indicate that MLKL knockdown by ASO does indeed impair necroptotic cell death *in vitro* similarly to what is observed in the atherosclerotic plaque *in vivo*.

### MLKL represses macrophage lipid loading

Because *Mlkl* knockdown reduced the formation of the necrotic core but did not decrease the total size of the atherosclerotic plaque, it suggests that the absence of MLKL promotes atherogenesis through a necroptosis-independent role. To further characterize the atherosclerotic lesions in MLKL ASO treated mice, we evaluated lesional foam cell accumulation using Oil Red O (ORO) staining for neutral lipids. We found that *Mlkl* silencing led to a significant increase in foam cell content within the plaque compared to controls (**Figure 3A**). Macrophage cholesterol balance is achieved through the processes of cholesterol uptake and removal pathways, therefore we first tested whether *Mlkl* knockdown inhibits the removal of cholesterol from plaque macrophages. We performed a cholesterol efflux assay in peritoneal macrophages after *Mlkl* knockdown, and found no differences in the efflux of radiolabelled cholesterol to either HDL or ApoA1 in MLKL ASO transfected macrophages (**Figure 3B**). To determine whether cholesterol uptake and retention, rather than efflux, was altered by loss of MLKL we incubated macrophages with cholesterol-rich lipoproteins acetylated, aggregated, or oxidized low-density lipoprotein (LDL). All forms of modified LDL treatment led to significantly higher levels of lipid accumulation in peritoneal macrophages after *Mlkl* knockdown (**Figure 3C**). Similar results were observed in bone marrow-derived macrophages (**Supplemental Figure 3A**). These findings were complemented by an increase in the uptake of radiolabelled cholesterol in peritoneal macrophages after *Mlkl* knockdown (**Supplemental Figure 3B**). These data together reveal a previously undefined role of MLKL in repressing lipid uptake and retention in macrophages that could act to limit the development of the atherosclerotic plaque.

**Figure 3.**
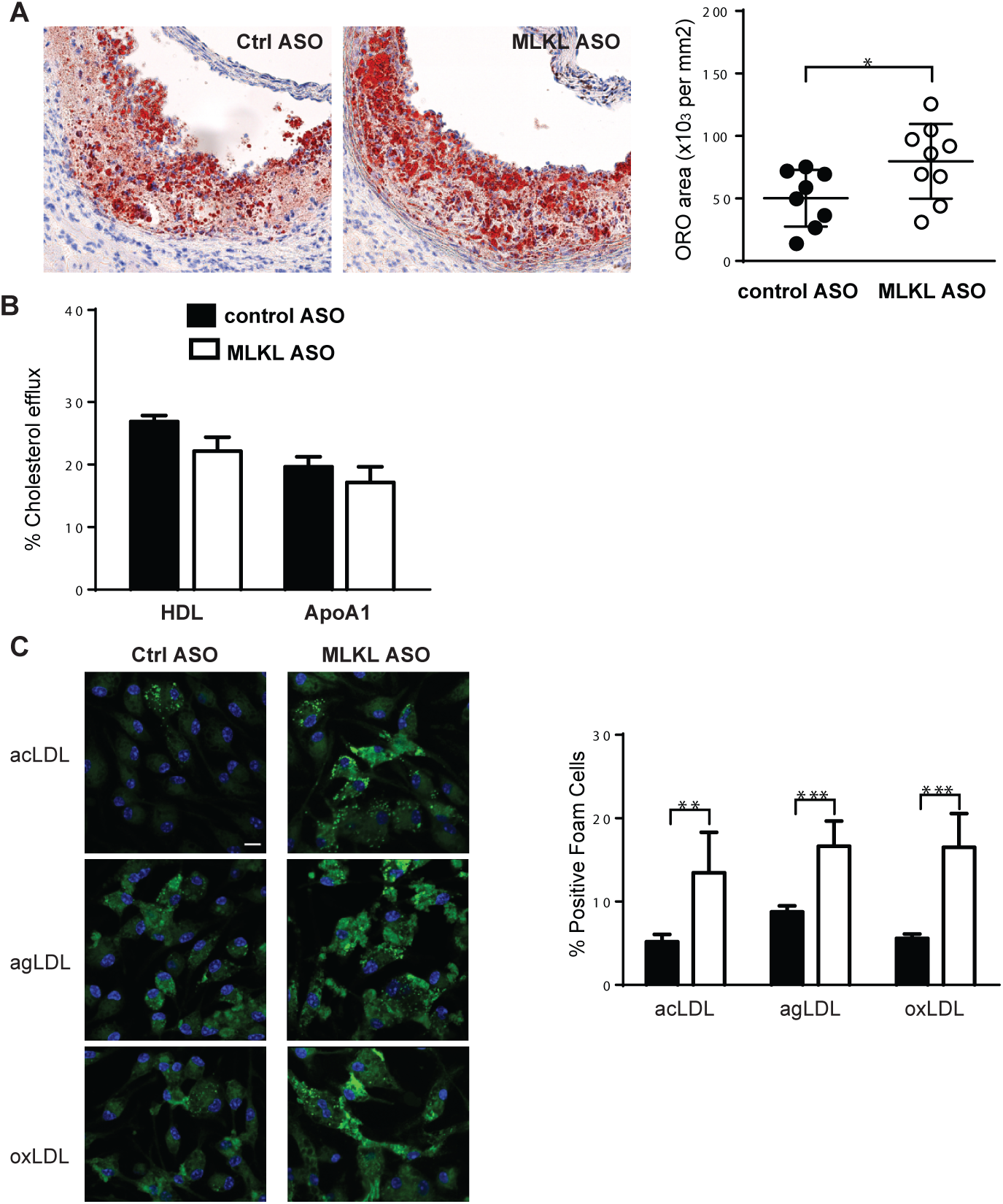
*Mlkl* knockdown increases the lipid content within the atherosclerotic plaque and isolated macrophages. **A**, Lipid content by Oil Red O staining was quantified in the aortic sinus plaques of *Apoe*-/- mice after 8 weeks of Western diet feeding and ASO treatment. Scale bar=200μm. Data presented as mean±SD. n=8-9 mice per group. **B**, Cholesterol efflux of radiolabelled cholesterol from peritoneal macrophages to HDL or ApoA1. **C**, BODIPY staining was performed in peritoneal macrophages after treatment with modified lipoproteins to assess proportion of foam-cell like macrophages. BODIPY: green; DAPI: blue. Scale bar=20μm. Data presented as mean±SEM. Representative experiments using n=5-7 mice per group. \*\**P*<0.01. acLDL: acetylated LDL; agLDL: aggregated LDL; oxLDL: oxidized LDL.

To better understand how MLKL alters cholesterol accumulation, we asked whether MLKL impacts the expression or activity of proteins involved in lipid uptake or efflux. We first assessed the expression of CD36 and ABCA1, a key scavenger receptor that mediates modified lipoprotein uptake and transporter of free cholesterol onto ApoA1 during efflux, respectively. Flow cytometry revealed no changes in the surface expression of ABCA1 or CD36 in MLKL ASO macrophages compared to controls (**Figure 4A**). Furthermore, western blotting revealed no change in ABCA1 or CD36 expression in these macrophages (**Supplemental Figure 4A**). Although MLKL does not have any kinase activity, it does interact with RIPK3, ESCRTIII and others, acting as a scaffold^18, 19^. As such, we examined the co-localization of endogenous MLKL with CD36 and ABCA1 by confocal microscopy. We did not observe any co-localization of MLKL with either CD36 (**Figure 4B, Supplemental Figure 4B**) or ABCA1 (**Figure 4C, Supplemental Figure 4C**). Although the uptake of modified lipoproteins proceeds though different receptors and pathways, they all share the common mechanism of endosomal trafficking and storage in lipid droplets. Therefore, we hypothesized that the increase in lipid loading with *Mlkl* knockdown was due to MLKL interaction with the endocytic trafficking machinery. To test this, we localized MLKL with early endosomal proteins EEA1 and Rab5, but did not observe any MLKL in these compartments under any of the lipid loading conditions tested (**Supplemental Figure 5A-B**). In contrast, we found colocalization of MLKL and the late endosomal marker Rab7, within large puncta in aggregated and oxidized LDL loaded macrophage (**Figure 4F**) but not acetylated LDL loaded macrophages (**Supplemental Figure 6**). These data therefore suggest that MLKL may restrict lipid loading in macrophages by regulating the endocytosis of modified lipoproteins within the macrophages.

**Figure 4.**
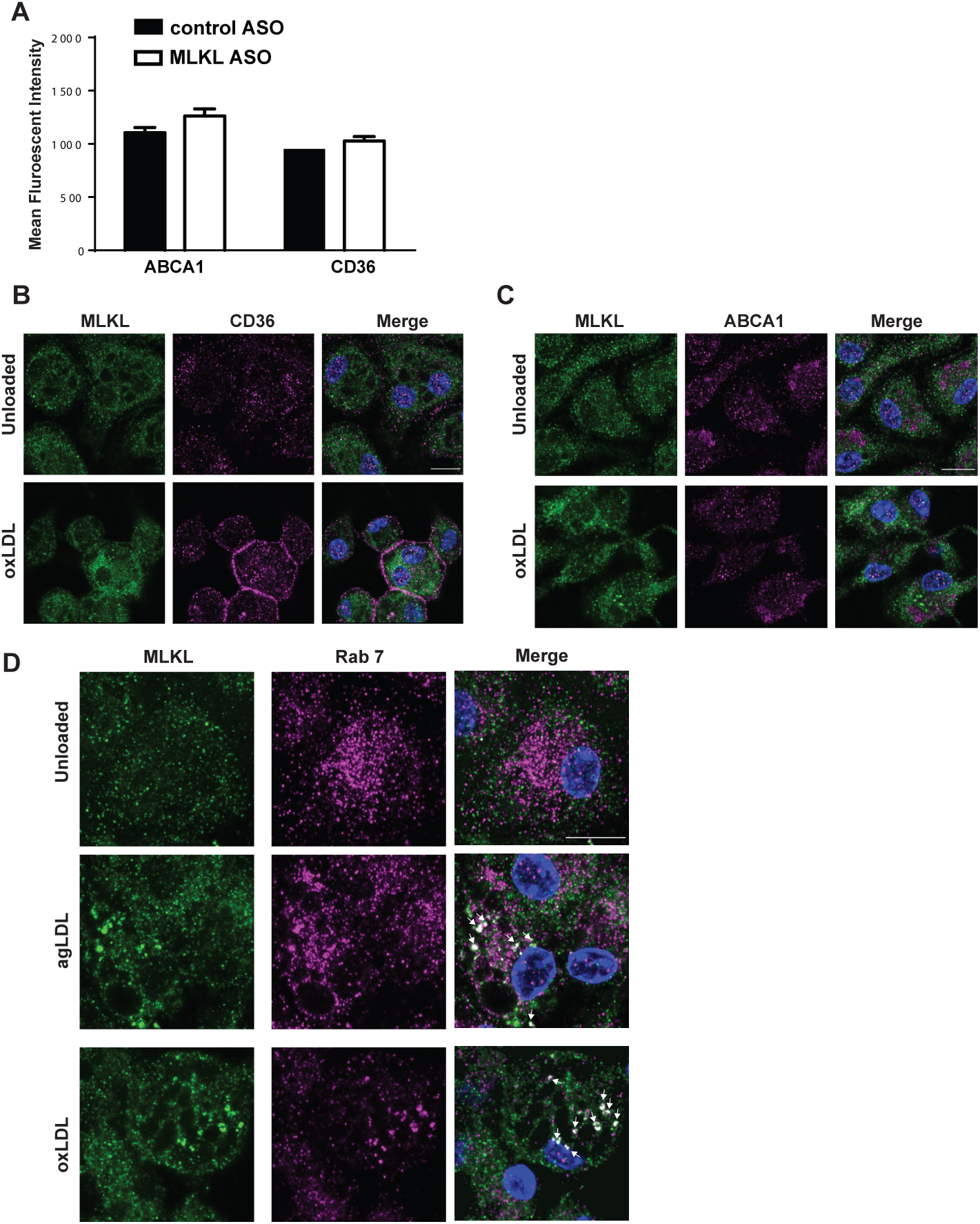
MLKL co-localizes with markers of late endosomes but not cholesterol receptors. **A**, Quantification of surface ABCA1 and CD36 were performed by flow cytometry analysis of peritoneal macrophages after transfection with ASOs and surface staining for CD36 or ABCA1. Data presented as mean±SEM. Representative experiments using n=4 mice per group. (**B-D**) Localization of MLKL with ABCA1 (B), CD36 (C) and Rab 7 (D) in peritoneal macrophages under control (unloaded) and or lipid loaded conditions. Co-localization is shown by white arrows. Scale bar=10μm. Representative images per condition.

## DISCUSSION

Necroptosis is a uniquely pro-inflammatory mode of cell death, initiated by the engagement of death receptors at the cell membrane, leading to the phosphorylation and oligomerization of MLKL and membrane lysis^5^. This mode of cell death plays prominent roles in cancer metastasis, inflammatory diseases, as well as in the normal immune response to pathogens^5, 20^. For this reason, inhibitors of the necroptosis signaling pathways are being investigated to mitigate the contributions from this mode of cell death to pathophysiologic processes. In our studies, we surmised that inhibition of the terminal protein involved in necroptosis, MLKL, would improve atherosclerosis through reduction of the necrotic core. In line with previous observations, where inhibition of other necroscome components reduced the advancement of the necrotic core^15, 16^, knockdown of *Mlkl* by antisense oligonucleotides also reduced formation of the necrotic core. However, unlike RIPK1 or RIPK3, this reduction in necrotic core did not reduce total lesion area. Furthermore, we observed an increase in lipid accumulation in the plaques of *Apoe*-/- mice receiving the MLKL ASO. In culture, isolated macrophages transfected with the MLKL ASO showed enhanced lipid loading upon incubation with modified lipoproteins, which was not due to defects in cholesterol removal from the cells. The majority of the previous studies using the *Mlkl*-knockout mice have performed studies with chow diet feeding, and focused on the role of MLKL in the necroptosis pathway. The studies outlined here are the first to assess chronic inhibition of MLKL in the context of high cholesterol diet feeding. Here we demonstrate that, in addition to its previously demonstrated function as the executioner of necroptosis, we have identified a previously unexpected role for MLKL in promoting retention of lipids in macrophages to drive atherogenesis. Together, these results indicate that MLKL regulates the cholesterol balance of macrophages independently from its role in necroptosis.

Upon *Mlkl* knockdown, we observed reductions in circulating cholesterol that were associated with increased lipid uptake in the plaque. These observations parallel what is seen in diseases where tissue macrophages accumulate lipids and plasma cholesterol levels are reduced^21, 22^. Our observations are also similar to the macrophage-specific knockouts of the cholesterol efflux transporters, *Abca1* and *Abcg1*, which also show increased macrophage lipid retention coupled with a decrease in circulating cholesterol level^23^. We therefore surmised that MLKL may interact with lipid efflux proteins to alter their activity. However, cholesterol efflux assays and Western blotting in peritoneal macrophages showed no changes in ABCA1 or CD36 protein levels or cell surface expression. This suggests that although the atherosclerosis phenotype in mice lacking *Mlkl* is similar to those lacking *Abca1*/*Abcg1*, the retention in lipids when MLKL is absent is likely via an ABCA1- and ABCG1-independent manner. Similarly, MLKL was not found to be localized at the membrane with either CD36, which is known to modulate oxidized LDL uptake, or ABCA1, which promotes free cholesterol efflux. It is noteworthy that these studies are among the first to evaluate endogenous MLKL expression and sub-cellular localization by microscopy in atherogenic macrophages.

MLKL has been shown to promote vesicle trafficking via interaction with the ESCRT machinery in the absence of pro-necroptotic stimuli^14, 19^. We therefore reasoned that MLKL may influence the endocytosis and trafficking of modified lipoproteins in foam cells. We found that while there was no co-localization with early endosome components EEA1 or Rab5, MLKL did co-localize with the late endosome marker Rab7. This interaction of MLKL and the late endosome may regulate the trafficking of cholesterol-rich lipoproteins, where the loss of MLKL prevents this trafficking and leads to increased lipid accumulation within the macrophages. The co-localization of MLKL and Rab7 was observed in distinct puncta of MLKL staining upon aggregated and oxidized LDL loading. These areas of co-localization may be oligomerized MLKL in its phosphorylated form, which we have shown previously is induced by oxidized LDL^15^ and has been previously described to regulate endosomal trafficking^14, 19^. However, MLKL and Rab7 were not co-localized under acetylated LDL conditions, which is known to be less inflammatory than oxidized and aggregated LDL, and where fewer MLKL positive punctate structures were seen. Others have shown *in vitro* that MLKL inhibits autophagic flux in non-macrophage cell types, therefore it is unlikely that the absence of MLKL would lead to reduced autophagy of lipid droplets^24,25^. Rather, we hypothesize that MLKL regulates the tethering of Rab7-positive late endosomes and/or multivesicular bodies (MVBs) at ER-endosome contact sites, possibly via ESCRT machinery to regulate lipid droplet mobilization^14, 19^. These distinct roles of MLKL in mediating necroptosis on the one hand and lipid droplet accumulation on the other likely explains why although necrotic core was reduced in the absence of MLKL, lipid accumulation was increased. Further studies are needed to fully delineate the role of MLKL in lipid droplet trafficking.

In conclusion, we find that MLKL participates in atherosclerosis by distinct mechanisms. Similar to the other necroptosis proteins, MLKL promotes lesion necrotic death, and loss of MLKL reduces necrotic core in advanced atherosclerotic plaques. However, loss of MLKL also promotes lipid droplet accumulation, likely as a result of its activity on endosomal trafficking, which acts to increase lipid retention in the plaque. These novel functions for MLKL in controlling cellular lipid metabolism suggest that MLKL likely plays multiple non-necroptotic roles in atherosclerosis and other metabolic diseases.

## ABBREVIATIONS

acLDL: acetylated low-density lipoprotein
agLDL: aggregated low-density lipoprotein
ApoA1: apolipoprotein A1
ASO: antisense oligonucleotide
DAMP: damage-associated molecular patterns
HDL: high-density lipoprotein
MLKL: mixed lineage kinase-like domain protein
ORO: Oil Red O
oxLDL: oxidized low density-lipoprotein
RIPK1: receptor interacting protein kinase 1
RIPK3: receptor interacting protein kinase 3

## ACKNOWLEDGMENTS

We would like to thank Xiaoling Zhao for her assistance with histology and Dr. James Murphy for his guidance with the MLKL microscopy.

## HIGHLIGHTS

- Similar to other necrosome components, MLKL expression drives necroptosis and the formation of the necrotic core during atherosclerosis.
- Independently from its role in cell death, MLKL elicits anti-atherogenic roles by limiting lipid accumulation in foam cells within the atherosclerotic plaque.
- MLKL colocalizes with the late endosome in macrophage foam cells, suggesting it may play a role in trafficking of internalized lipids to mediate their accumulation.

## MATERIALS AND METHODS

### Mice

All animal procedures were approved by the University of Ottawa Animal Care and Use Committee. Male and female WT and *Apoe*-knockout (*Apoe*-/-) mice were maintained on a chow diet (#2016, Harlan Teklad) in a temperature and light-controlled environment. For the atherosclerosis studies, 8-9-week old *Apoe*-/- mice were fed a Western style diet (TD 88137; 21.2% fat, 0.2% cholesterol; Harlan Teklad) for 8 or 16 weeks, while receiving weekly subcutaneous injections of 50mg/kg control or two independent MLKL-targeting antisense oligonucleotides (ASOs, Ionis Pharmaceuticals). The *Apoe*-/- mice were fasted prior to sacrifice. The mice were euthanized under isoflurane anesthesia by exsanguination. Aortic sinuses were immediately embedded in OCT (Cedarlane) and frozen until processing.

### Aortic Sinus Histology

Aortic sinuses were cryosectioned using a Leica CM 3050S cryostat at a thickness of 10μm and serial sections were placed on Superfrost Plus microscope slides (Fisher Scientific). The sections were stored at −20°C until use. Lesions were identified by hematoxylin & eosin (H&E) staining. Cell death in the aortic sinus was assessed by TUNEL staining kit (Roche) following the manufacturer’s recommendations. Oil Red O staining of the aortic sinus sections were performed at 60°C for 12 minutes. Images were acquired using a Leica Aperio 8 slide scanner or Zeiss LSM 880 confocal microscope.

### Serum collection, cholesterol quantification, and multiplex ELISA analyses

At sacrifice, whole blood was collected in serum collection tubes (Becton Dickinson) and was centrifuged at 10000*x*g for 2 minutes for the separation of serum. Cholesterol was measured in frozen serum by enzymatic assay (Cholesterol E Kit, Wako). Additionally, frozen serum was used for the quantification of cytokines using the Pro-inflammatory Cytokine 23-plex Bio-plex kit (Bio-Rad) according to manufacturer’s recommendations.

### Complete blood counts

Whole blood at sacrifice was collected in EDTA-coated microvettes. The samples were then analyzed on a Hemavet 950 (Drew Scientific).

### Isolation and transfection of thioglycolate elicited peritoneal macrophages

Peritoneal macrophages were collected from WT mice 4 days after intraperitoneal injection of 1ml thioglycolate. Control or one of two MLKL ASOs (50mg/kg) were injected into the peritoneal cavity 30-36 hours prior to harvesting. The cells were seeded in 10% FBS/DMEM overnight prior to experiments. Results for both MLKL-targeting sequences were pooled for analyses.

### Western blot analysis

Samples were lysed in RIPA buffer containing protease and phosphatase inhibitors (Roche). Proteins were subjected to SDS-PAGE and then transferred to PVDF membranes (Bio-Rad). Immunoblotting was performed for MLKL (Millipore), GAPDH (Millipore), and ABCA1 (Novus Biologicals). Horseradish peroxidase-conjugated secondary antibodies (GE Healthcare) and ECL Clarity or Clarity Max (Bio-Rad) were used to detect proteins on a ChemiDoc XRS+ (Bio-Rad).

### Cell viability assay

Cell death after treatment with necroptotic stimuli was assessed as previously described^15^. Here, peritoneal macrophages were treated with 50ng/ml TNFα (Thermo Fisher Scientific), 50μM zVAD.fmk (APExBio), and 50μM necrostatin-1 (Sigma-Aldrich) in 2% FBS/DMEM for 6 hours prior to harvesting the cell culture media.

### Cholesterol efflux

Cholesterol efflux from peritoneal macrophages was assessed as previously described^26^. Briefly, macrophages were incubated with radiolabelled cholesterol (Perkin Elmer) incorporated into acetylated low-density lipoproteins for 24 hours, followed by an overnight incubation period in 0.2% fatty acid free albumin (Sigma-Aldrich), and efflux was performed upon incubation with either HDL (Alfa Aesar) or recombinant human ApoA1 for an additional 24 hours.

### Fluorescence microscopy

Modified lipoproteins were generated as previously described^26^. Peritoneal macrophages were loaded with 100μg/ml of acetylated, oxidized, or 50μg/ml aggregated low-density lipoproteins for 30 hours, followed by an overnight equilibration period in 0.2% fatty acid free albumin. Macrophages were fixed in 4% paraformaldehyde and immunofluorescence staining was performed for neutral lipids (BODIPY 493/503, Thermo Fisher Scientific). Images were acquired using an LSM 880 Zeiss confocal microscope with Airyscan mode. Macrophages were manually scored as lipid-loaded by the presence of lipid droplets or reported as percent of BODIPY positive area. For immunofluorescent staining, macrophages loaded with 100μg/ml of acetylated, oxidized, or 50μg/ml aggregated low-density lipoproteins for 30 hours then were fixed in ice cold methanol and stained with MLKL (Millipore, 1:50), ABCA1 (Novus Biologicals), CD36 (Novus Biologicals), EEA1 (Abcam, 1:200), Rab5 (Abcam 1:1000), and Rab7 (Abcam, 1:500) using goat anti-rat Alexa Fluor 555 (1:500) or donkey anti-rabbit Alexa Fluor Plus 647(1:1000, both,Thermo Fisher Scientific).

### Flow cytometry

Macrophages were incubated with ABCA1 (Novus Biologicals) or CD36 (Novus Biologicals) followed by goat anti-rabbit Alexa Fluor 594 secondary antibodies, all at a 1:100 dilution. Samples were acquired on a FACS Aria IIIu (BD Biosciencces) flow cytometer.

### RNA expression and gene expression analysis

Livers were homogenized and total RNA was isolated using Trizol (Invitrogen). Reverse transcription and gene expression analysis was performed as previously described^27^. Genes of interest were normalized to mHprt. The primers used are as follows: mHprt (Forward: 5’ TCAGTCAACGGGGGACATAAA 3’; Reverse: 5’ GGGGCTGTACTGCTTAACCAG 3’) and mMlkl (Forward: 5’ AATTGTACTCTGGGAAATTGCCA 3’; Reverse: 5’ TCTCCAAGATTCCGTCCACAG 3’).

### FPLC cholesterol quantification

Fresh serum was used for separation of lipoproteins by FPLC as previously described^28^. Cholesterol was quantified in fractions from fresh pooled serum by enzymatic assay (Cholesterol E Kit, Wako).

### Bone marrow derived macrophage isolation and microscopy

Bone marrow was isolated from femurs and tibiae of WT mice fed a Western diet and administered control or MLKL ASO for 15 weeks. The bone marrow was cultured in 20% L929/10% FBS/DMEM media for 7 days prior to lipid loading with 50μg/ml acetylated or 100μg/ml aggregated low-density lipoproteins (Alfa Aesar) as described previously. Images were acquired on a BioTek Cytation5 Cell Imaging Multi-Mode Reader.

