## Supplemental Figure for "Loss of MLKL Decreases Necrotic Core but Increases Macrophage Lipid Accumulation In Atherosclerosis"

#### Supplemental Figure 1

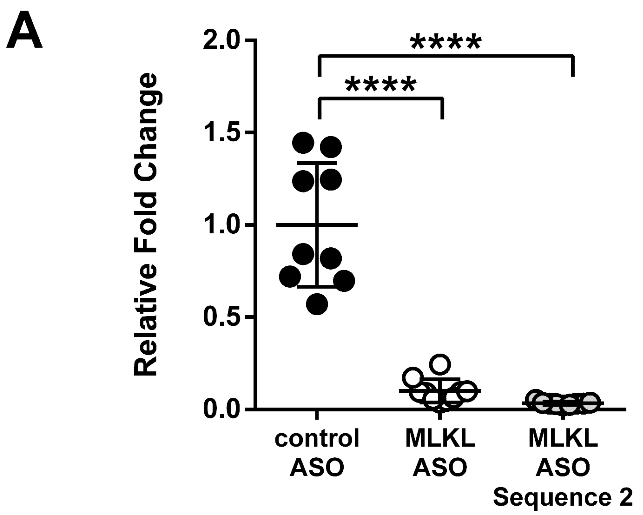


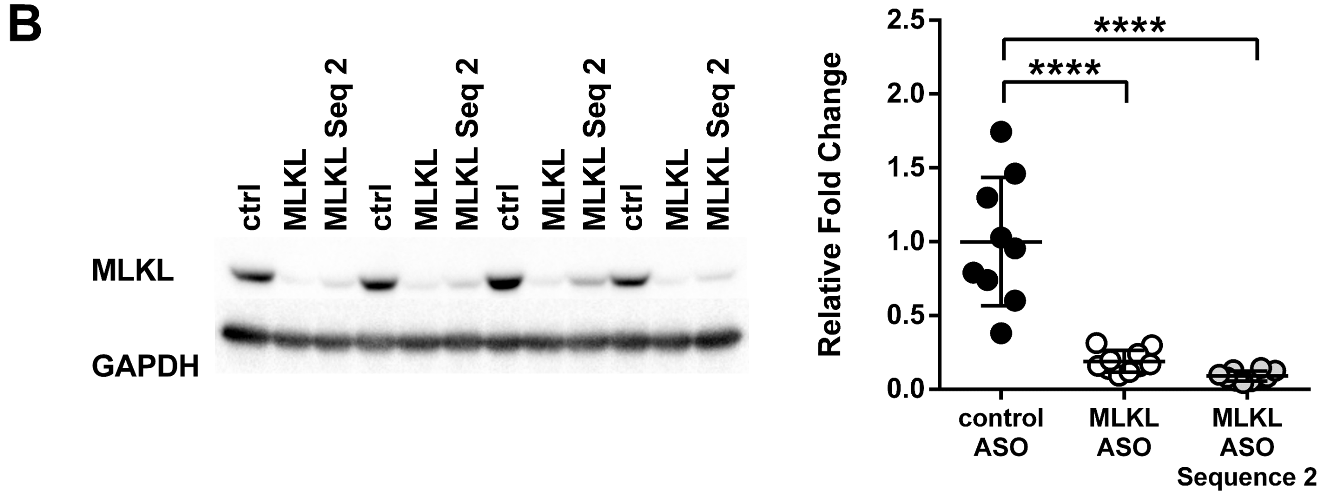


**Supplemental Figure 1.** Treatment with MLKL ASOs efficiently knocks down Mlkl both at the gene and protein expression levels. Livers were isolated from the *Apoe*-/- mice after 8 weeks of diet and ASO treatment where **A,** gene expression and **B,** Western blotting was performed to confirm Mlkl knockdown. Data presented as mean±SD. n=9-10 mice per group. ****$P$<0.0001.

### Supplemental Figure 2

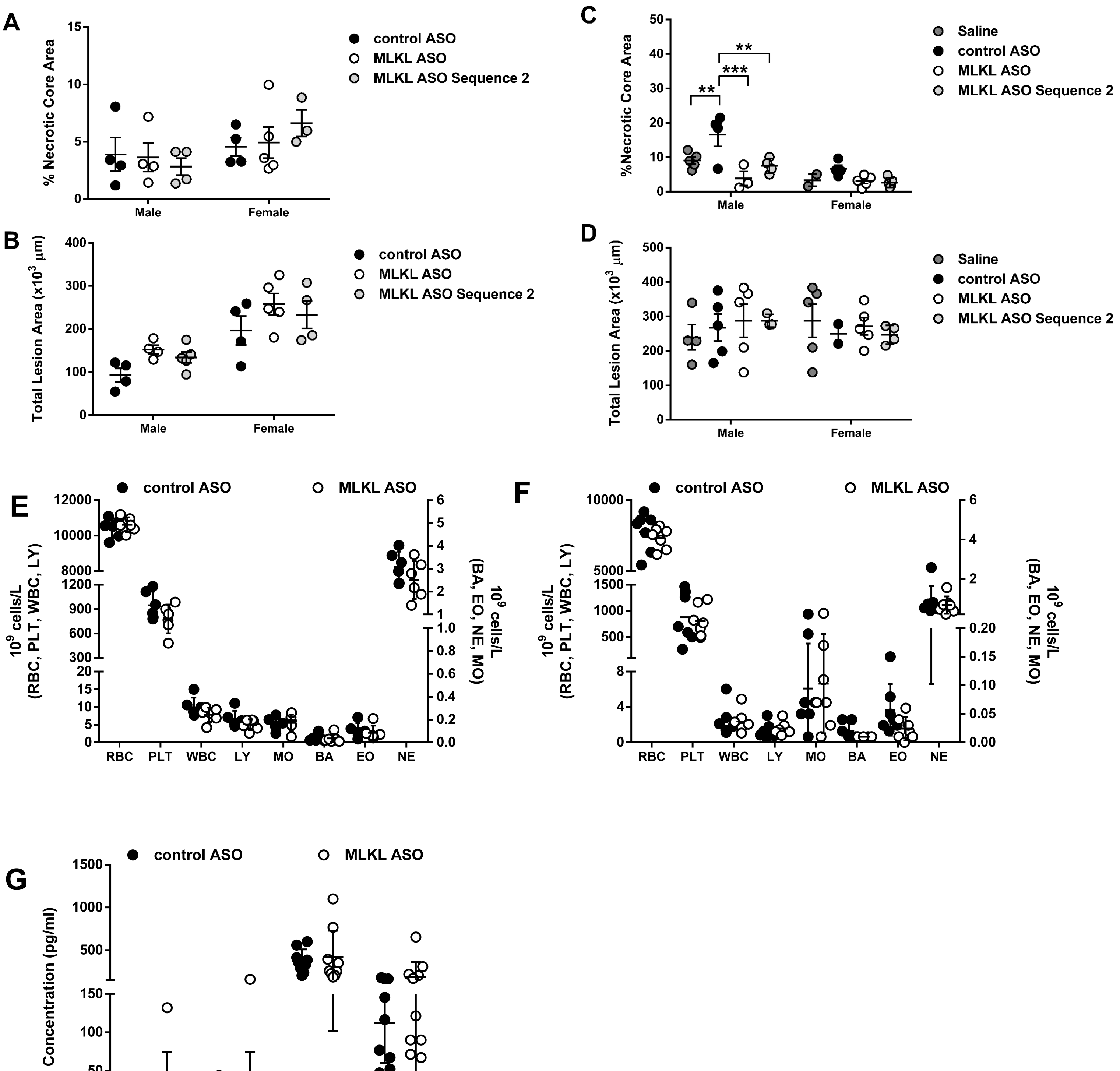


**Supplemental Figure 2.** Total and necrotic core areas after treatment with MLKL ASOs. Total lesion area and percent necrotic core areas were assessed in the aortic sinus of *Apoe*-/- mice fed a Western diet and treated with ASOs for 8 weeks (**A-B**) and 16 weeks (**C-D**). Data presented as mean±SD. n=2-5 mice per group. **E-F**. Complete blood counts were performed in whole blood after **E,** 8 and **F,** 16 weeks of treatment. **G,** Circulating pro-inflammatory cytokines were quantified by multiplex ELISA in the serum of *Apoe*-/- mice treated for 16 weeks (select factors shown). n=6-11 mice per group. Data presented as mean±SD.
**$P$<0.01, ***$P$<0.001.

### Supplemental Figure 3

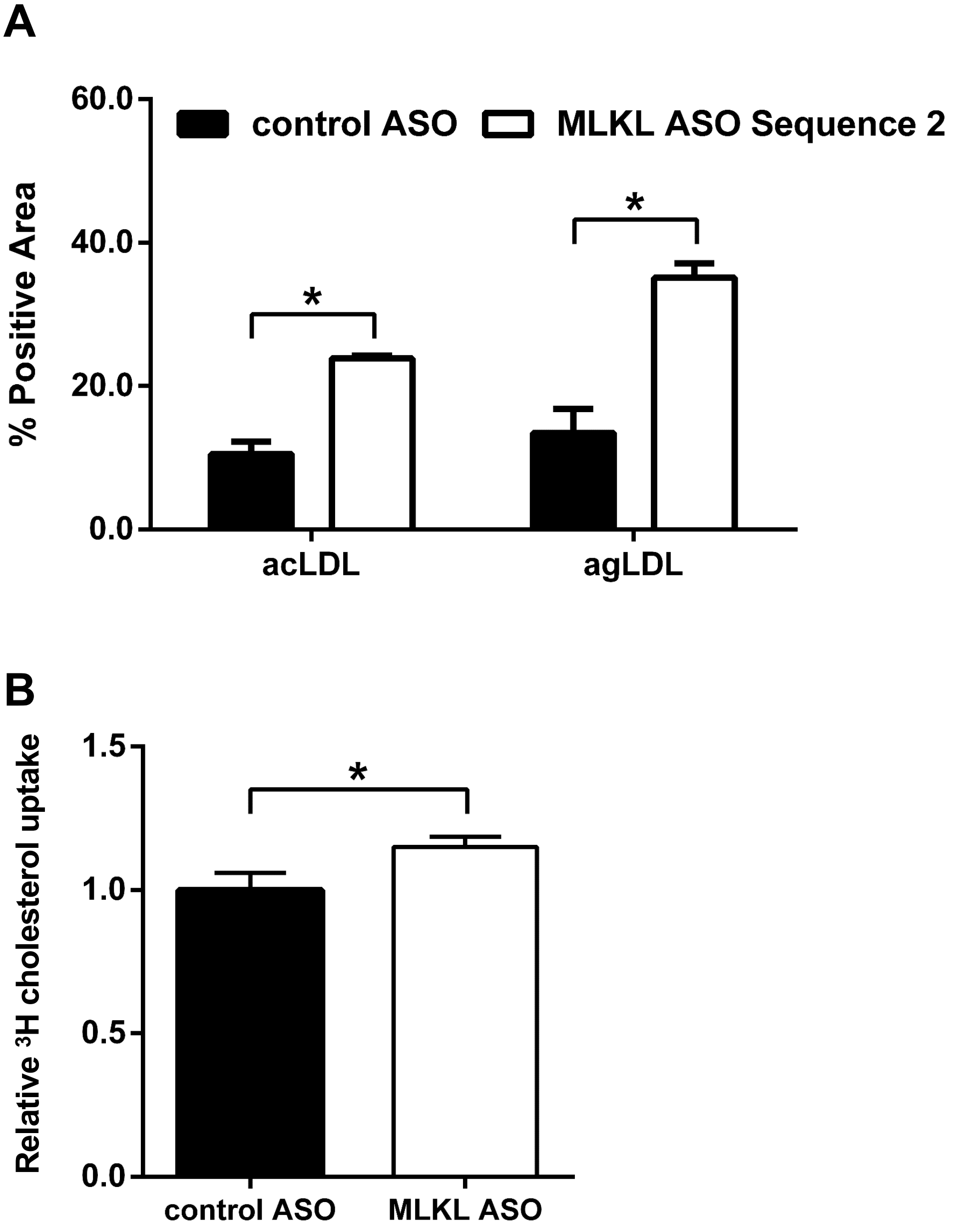


**Supplemental Figure 3.** Mlkl knockdown enhances macrophage lipid loading. **A,** Bone marrow derived macrophages were isolated from WT mice treated with control or MLKL ASO sequence 2 and lipid loaded with acetylated or aggregated low-density lipoprotein. BODIPY staining was used to quantify intracellular lipids. Representative experiment using 2 mice per group. **B**, Cholesterol loading using radiolabelled cholesterol in peritoneal macrophages was reported as fold change compared to control ASO.. Data presented as mean±SEM. *$P$<0.05.

Supplemental Figure 4

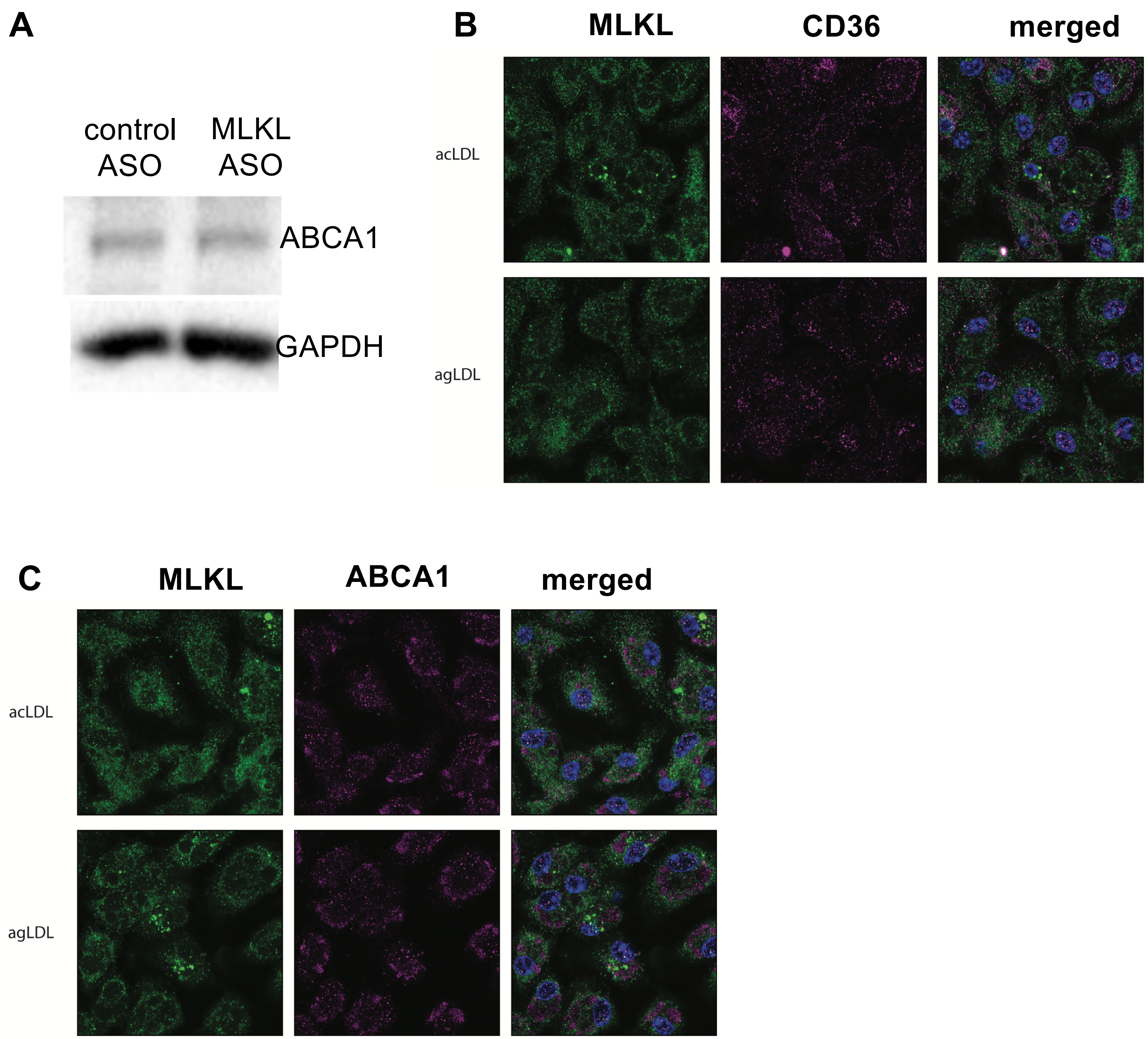


**Supplemental Figure 4. A,** Immunoblotting was performed for Abca1 in peritoneal macrophages transfected with ASOs. Representative experiment using 5-11 mice per group. **B-C.** Colocalization of MLKL with CD36 (B) or ABCA1 (C) was performed in peritoneal macrophages under control and acetylated and aggregated LDL loaded conditions. Scale bar=10μm. Representative images per condition.

### Supplemental Figure 5

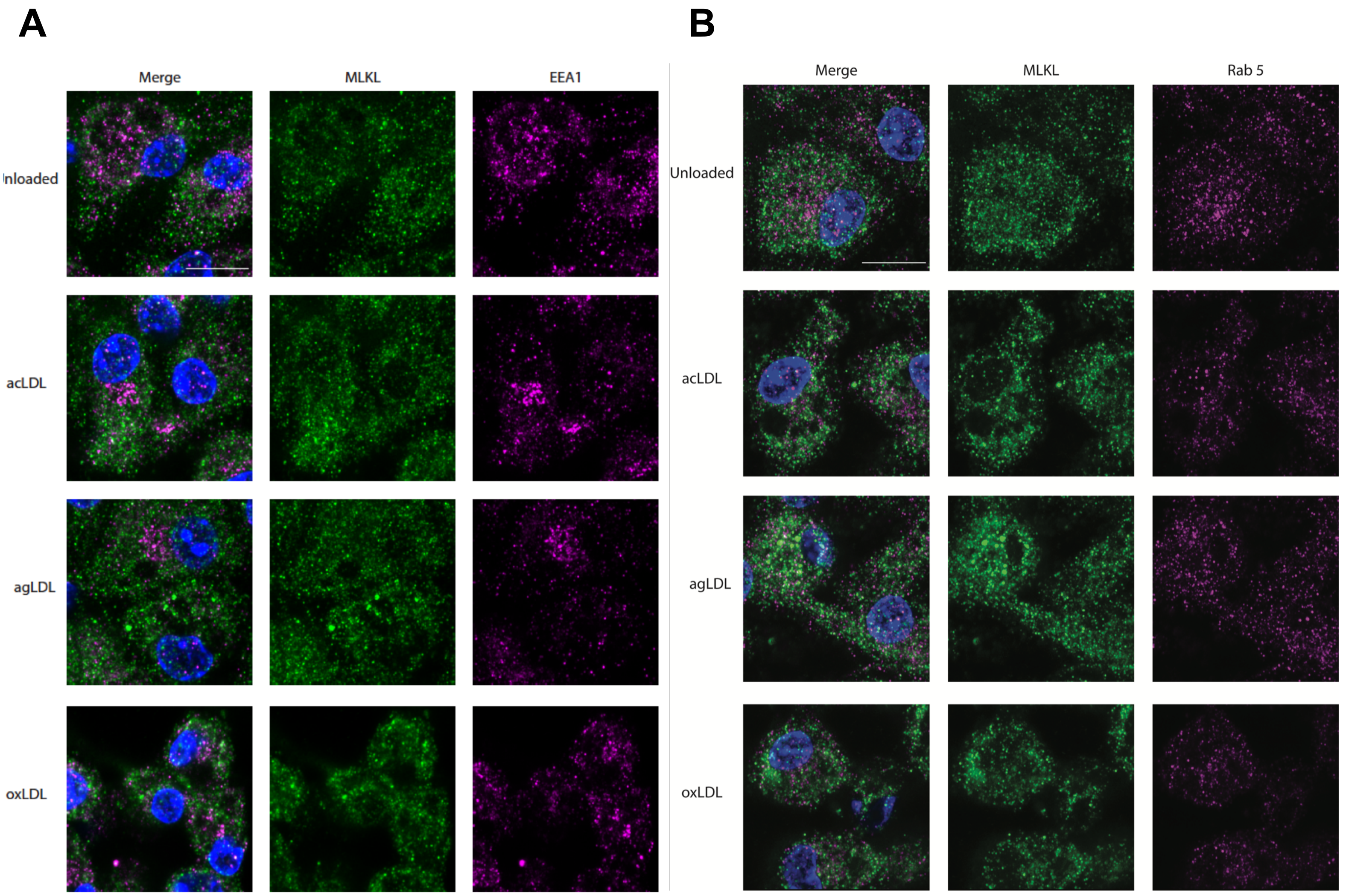


**Supplemental Figure 5.** Localization of MLKL with EEA1 (A) or Rab5 (B) in peritoneal macrophages under control, acetylated, aggregated or oxidized LDL loaded conditions. Scale bar=20μm. Representative images per condition.

### Supplemental Figure 6

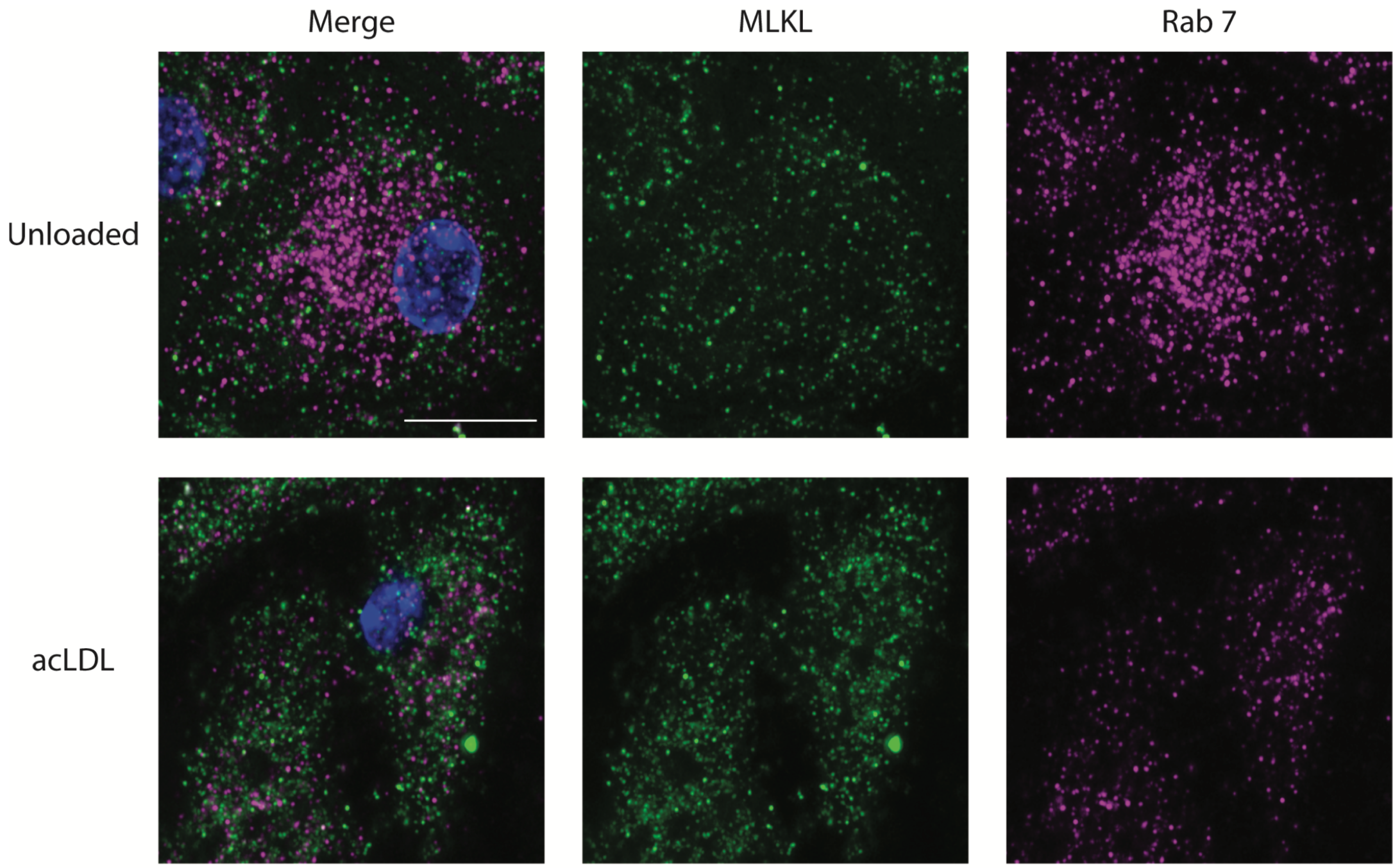


**Supplemental Figure 6.** **L**ocalization of MLKL and Rab7 in peritoneal macrophages under control or acetylated LDL loaded conditions. Scale bar=20μm. Representative images per condition.
